# NicheAgent: LLM-Guided Zero-Shot Niche Identification for Spatial Transcriptomics

**DOI:** 10.64898/2025.12.09.693287

**Authors:** Sajib Acharjee Dip, Liqing Zhang

## Abstract

Spatial transcriptomics provides high-resolution maps of gene expression within intact tissue architecture, enabling the study of cellular niches, functional layers, and microenvironmental structure. Yet, accurately assigning niche or layer identities remains challenging across platforms such as 10x Visium, MERFISH, and STARmap due to batch variability, incomplete marker panels, and the lack of universally consistent domain boundaries. Existing methods including SpaGCN, BayesSpace, STAGATE, and DeepST rely heavily on supervised labels, dataset-specific fine-tuning, or deep representation learning, which often over-smooth boundaries, fail to generalize across technologies, and provide limited interpretability.

We introduce **NicheAgent**, a zero-shot, training-free framework for spatial niche identification guided by lightweight large language models (LLMs). NicheAgent constructs interpretable *nichecards* for each region, encoding canonical marker genes and prototype expression centroids. Each cell is first assigned using a deterministic nearest-prototype rule based solely on gene expression and 2-hop spatial neighborhoods. Low-confidence assignments are then selectively reviewed and corrected by an LLM using only interpretable signals: marker-gene coherence, neighborhood label consistency, and an allowed label set. A final spatial smoothing step enforces local structural coherence.

Applied across Visium, MERFISH, and STARmap tissues without any retraining or domain-specific supervision, NicheAgent achieves robust and biologically meaningful niche delineation, outperforming many supervised and graph-based baselines on homogeneity, completeness, and mutual information, while offering transparent reasoning traces for each corrected decision. Our results demonstrate that LLM-guided refinement, when coupled with lightweight rule-based prototypes, provides a scalable, explainable, and cross-platform alternative to heavy deep learning models for spatial transcriptomics annotation.

## 1 Introduction

Spatial transcriptomics (ST) technologiesincluding MERFISH [3], STARmap [19], and 10x Visium enable in situ measurement of gene expression while preserving the native spatial organization of tissues. These technologies have revealed fine-grained anatomical structure, functional compartments, and microenvironmental organization across diverse tissues and subjects. However, accurately annotating spatial regions (e.g., cortical layers, cellular niches, or anatomical domains) remains challenging due to several factors: heterogeneous gene coverage across platforms, variable spatial resolution, cross-subject biological variability, inconsistent marker robustness, and the absence of matched ground-truth annotations for many experiments.

Early computational approaches such as Giotto [9] and SpatialDE [17] primarily relied on spatial statistics and unsupervised clustering based on expression covariance. Later graph-based models including BayesSpace [23], SpaGCN [11], STAGATE [7], and GraphST [13] combined expression with spatial neighborhood graphs to encourage local smoothness. Although effective on many datasets, these methods often require extensive hyperparameter tuning, may blur sharp anatomical boundaries, and typically do not incorporate prior biological knowledge beyond learned embeddings. Their interpretability also remains limited, especially in cases where marker gene signals conflict with spatial proximity or when datasets contain sparse or technically noisy gene panels (as commonly seen across multiple subjects in MERFISH and STARmap).

Large language model (LLM) driven annotation methods have recently emerged, using natural-language reasoning over gene sets or functional signatures, as seen in models such as scGPT [5] and GenePT [4]. However, these approaches rely heavily on textual prompts, require substantial model capacity, and do not directly operate on the structured numerical nature of ST data. Furthermore, most LLM workflows treat annotation as a single-step decision, lacking an explicit arbitration mechanism between quantitative signals (e.g., prototype distances) and contextual biological cues (e.g., neighborhood marker enrichment).

Motivated by how human experts annotate spatial data first applying simple, well-understood heuristics, then revising ambiguous regions by integrating marker genes, local neighborhoods, and prior anatomical knowledge we introduce **NicheAgent**, a lightweight, zero-shot annotation framework designed for heterogeneous ST datasets across multiple subjects and platforms (Visium, MER-FISH, and STARmap). Instead of training new models, NicheAgent combines three complementary components: (1) *prototype-based rule assignment*, using curated region centroids and canonical marker sets; (2) a *constrained LLM reviewer* that is invoked only for low-confidence cells and restricted to a predefined region ontology; and (3) *one-round spatial smoothing* to enforce anatomical coherence while avoiding the over-smoothing characteristic of graph-based deep models.

Unlike recent transformer-based spatial architectures such as DeepST [21], SpaFormer [20], and Sopa [1], which primarily focus on learning improved spatial representations, NicheAgent emphasizes transparency, interpretability, and minimal parameterization. The method is inherently robust across subjects and data modalities because it relies on structured biological knowledge, local marker enrichment, and constrained reasoning rather than large-scale model training or extensive hyperparameter search.

Across multiple datasets spanning MERFISH, STARmap, and Visium with several subjects per technology, we find that simple prototype-based heuristics remain strong baselines, while the LLM reviewer improves annotation quality in ambiguous regions, particularly where marker signatures are noisy or partially missing. These results suggest that small, ontology-constrained LLMs can serve as effective and computationally efficient “reviewers,” complementing traditional model-based heuristics rather than replacing them entirely.

## 2 Methods

We introduce **NicheAgent**, a zero-shot spatial annotation framework that integrates (i) prototype-based region inference using curated *NicheCards*, (ii) neighborhood-level marker enrichment, and (iii) a constrained large-language-model (LLM) reviewer for difficult cases (shown in Figure 1). Unlike supervised spatial models [9, 8, 18], NicheAgent requires *no* training labels and instead leverages bio-logical priors and region prototypes derived from external datasets.

**Fig. 1.**
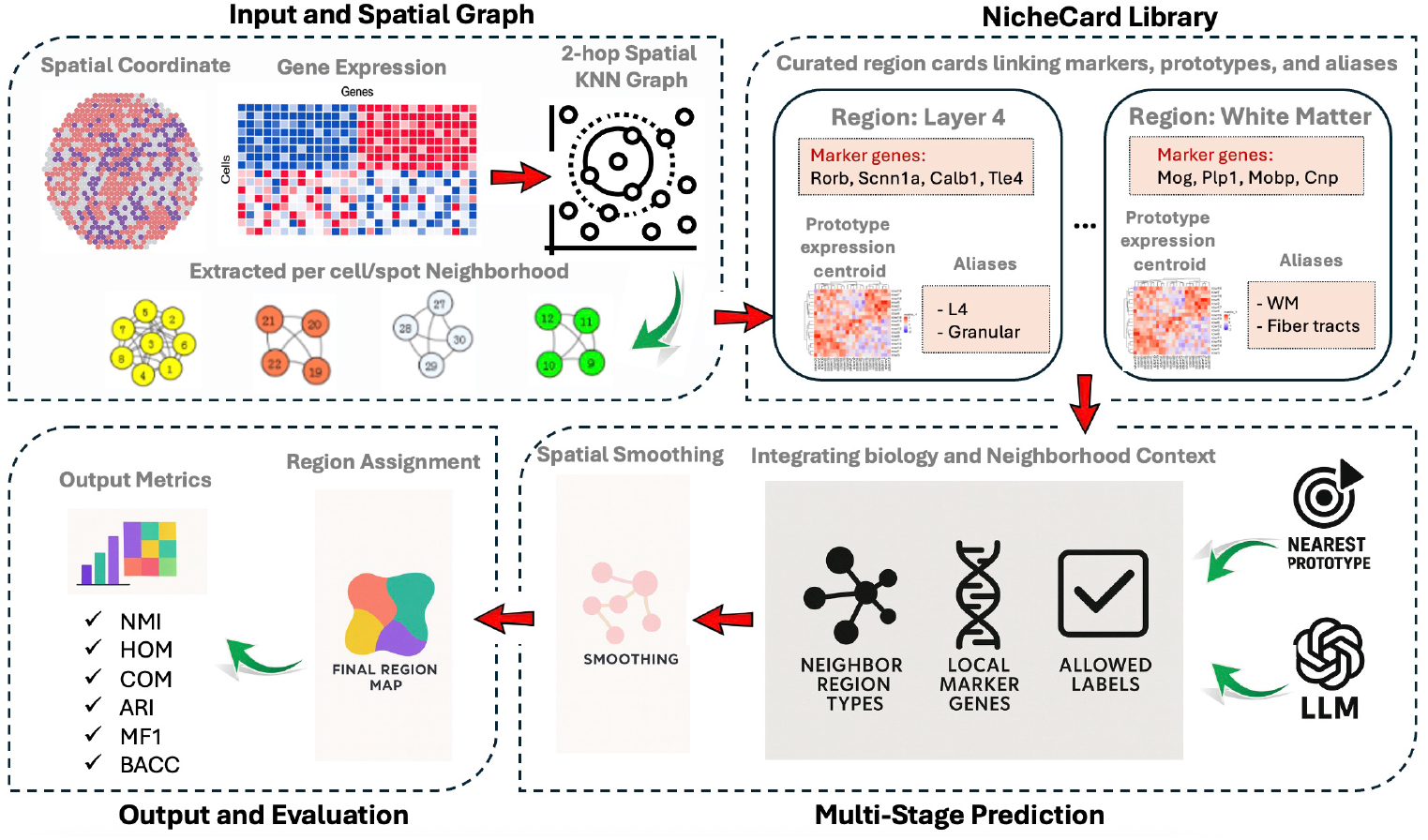
Overview of the NicheAgent framework for zero-shot spatial niche annotation. NicheAgent operates in three stages. (a) Input and Spatial Graph: For each tissue, spatial coordinates and gene-expression matrices are used to build a 2-hop spatial k-nearest-neighbor (KNN) graph, extracting neighborhood-level gene and spatial features per cell or spot. (b) NicheCard Library: Curated region prototypes (“nichecards”) link canonical marker genes, prototype expression centroids, and human-readable aliases (e.g., Layer 4 – Granular, White Matter – Fiber tracts). (c) Multi-stage Prediction: Each cell is first assigned to its nearest prototype using deterministic rules based on local marker-gene coherence and neighbor-type statistics. Low-confidence assignments are selectively reviewed and refined by a lightweight large language model (LLM) that reasons over allowed labels and biological context, producing interpretable rationales for each correction. A final spatial-smoothing step enforces local coherence within the tissue architecture. (d) Output and Evaluation: The resulting region map is evaluated by standard clustering and segmentation metrics (NMI, HOM, COM, ARI, macro-F1, balanced accuracy).

### 2.1 Notation

Let *X* ∈ ℝ^*n*×*p*^ denote the gene expression matrix of *n* cells/spots and *p* genes. Let *s*_*i*_ ∈ ℝ^2^ denote the spatial coordinates of cell *i*. Let = {1, …, ℛ} denote the set of anatomical regions. For each region *r*, let *µ*_*r*_ ∈ ℝ^*p*^ denote its prototype centroid and *M*_*r*_ its curated canonical marker set. Let *G* = (*V, E*) denote the spatial graph with node set *V* = {1, …, *n*} and edges defined via radius-based proximity.

The objective is to infer for each cell *i* a label

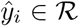

without using annotated spatial transcriptomics data.

### 2.3 Input Data and Spatial Graph Construction

Given an AnnData object containing gene expression and spatial coordinates, NicheAgent constructs a 2-hop radius-based spatial graph. Two cells *i, j* are adjacent if:

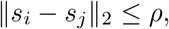

where *ρ* is dataset-dependent (e.g., *ρ* = 50 for Visium, *ρ* = 250 for STARmap).

Define the first-hop neighborhood:

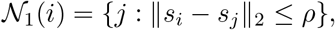

and the 2-hop adjacency:

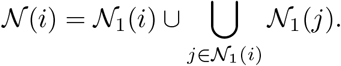

This 2-hop expansion improves robustness to sparse regions and noisy boundaries.

#### Algorithm 1

BuildSpatialGraph: Radius-Based 2-Hop Graph Construction

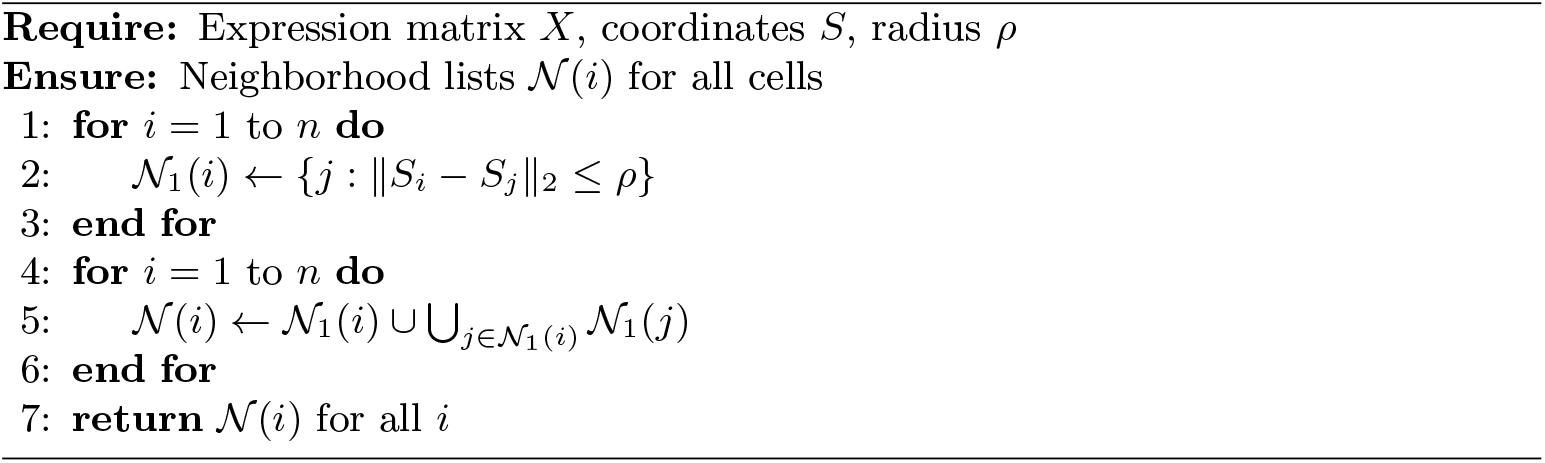

### 2.3 Curated NicheCards: Region Prototypes and Marker Gene Priors

Each region *r* is represented by a **NicheCard**:

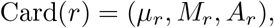

where *µ*_*r*_ is the prototype centroid computed from external high-confidence single-cell or spatial datasets, *M*_*r*_ is a curated canonical marker set collected from reference atlases [22, 10], and *A*_*r*_ is an alias/synonym set used for robust LLM-based matching.

These prototypes act as biologically grounded anchors enabling zero-shot inference, consistent with prototype-based learning paradigms [16].

### 2.4 Local Marker Gene Extraction

For each cell *i*, the neighborhood-enriched marker score for gene *g* is:

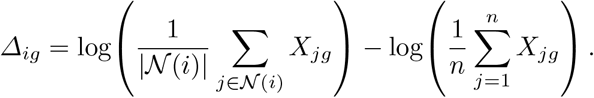

We select the top-*k* genes by *Δ*_*ig*_ (default *k* = 30), forming the set

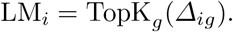

These genes help the LLM infer the biological identity of difficult cells.

### 2.5 Prototype-Based Zero-Shot Label Assignment

For each cell *i*, we compute the squared Euclidean distance to each region prototype:

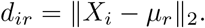

Let:

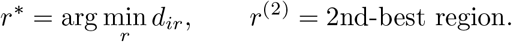

#### Algorithm 2

RuleBasedInference: Nearest-Prototype Assignment

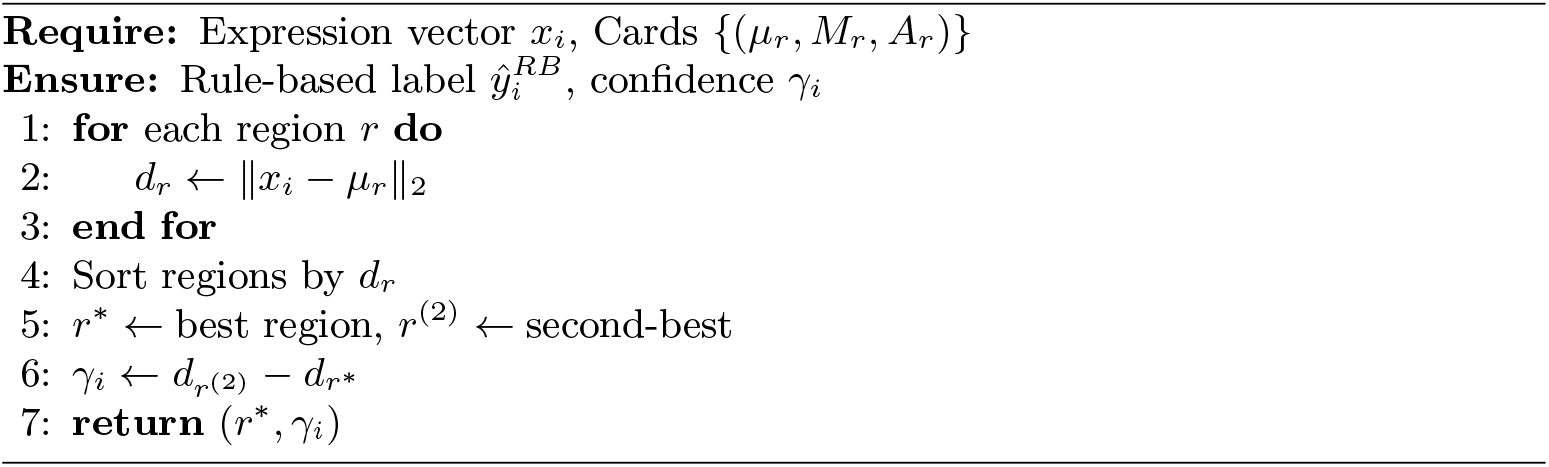

Define a confidence margin:

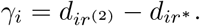

A large *γ*_*i*_ indicates a confident match. Cells with small margin or low neighborhood coherence are escalated to the LLM reviewer.

### 2.6 Low-Confidence Cases: Constrained LLM Reviewer

We use the Qwen-2.5 14B model (Ollama backend, deterministic decoding with temperature = 0) as the constrained LLM reviewer for adjudicating low-confidence cells. For ambiguous cells (*γ*_*i*_ ≤ *τ*), we query a deterministic LLM [2, 14, 6] with a structured, biology-informed prompt containing:

1. the rule-based proposed label *r*^∗^,
2. the allowed region label set,
3. neighbor region-type frequencies,
4. local marker genes LM_*i*_.

The LLM returns:

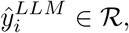

parsed using strict JSON and cleaned with alias sets *A*_*r*_.

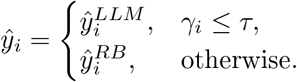

### 2.7 Spatial Smoothing

Following LLM adjudication, we apply a single round of spatial smoothing:

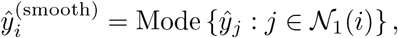

with ties resolved by keeping 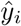. One round prevents oversmoothing while removing salt-and-pepper noise.

#### Algorithm 3

LLMReview: Constrained Label Adjudication

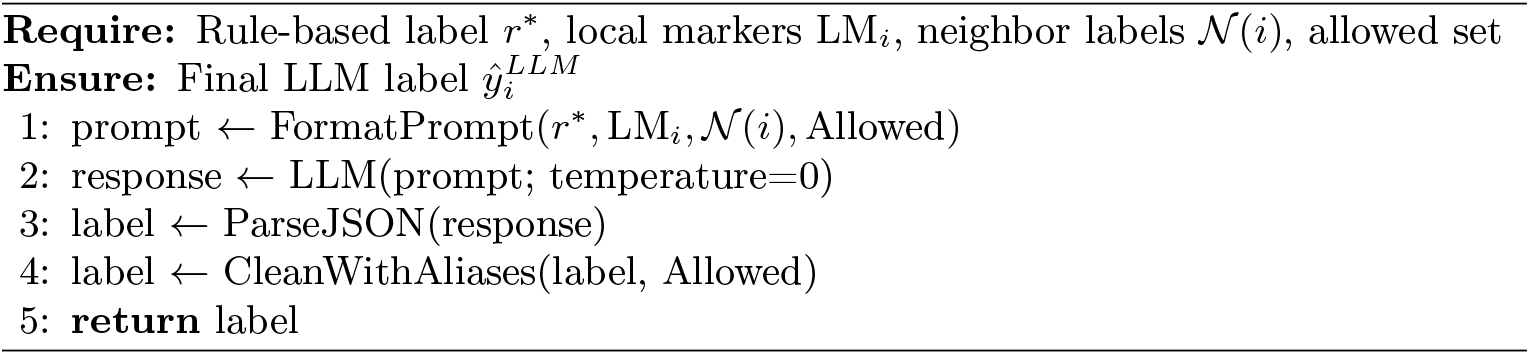

### 2.8 Evaluation Metrics

We compute standard clustering/annotation metrics, including NMI, homogeneity, completeness [15], ARI [12], accuracy, balanced accuracy, macro-F1, and silhouette score in 50-dimensional PCA space.

NMI uses the sklearn normalized formulation:

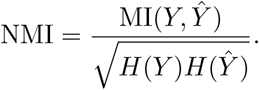

### 2.9 Computational Complexity

Let *n* be the number of cells and *R* the number of regions.

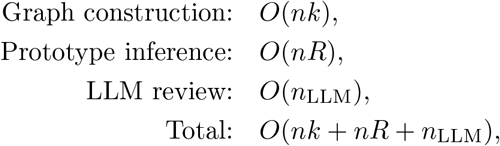

where *n*_LLM_ ≪ *n* because only low-confidence cells are escalated.

## 3 Results

### 3.1 Performance Across Visium, MERFISH, and STARmap Datasets

Across all three datasets and all evaluation metrics (NMI, HOM, and COM), **NicheAgent** consistently achieves the strongest performance, outperforming both classical baselines and recent LLM-based annotators.

**First**, NicheAgent attains the highest average NMI (**0.8045**) across Visium, MERFISH, and STARmap, substantially exceeding the top prior LLM-based baseline (LLMiniST-Fs: 0.686) as shown in Table 2. The gains are most prominent on MERFISH, where NicheAgent reaches **0.9208** ± **0.015**, representing a 10–15% improvement over the next best method. This indicates that the combination of prototype-based assignment and targeted LLM adjudication yields highly accurate region recovery without any training (Figure 2).

**Fig. 2.**
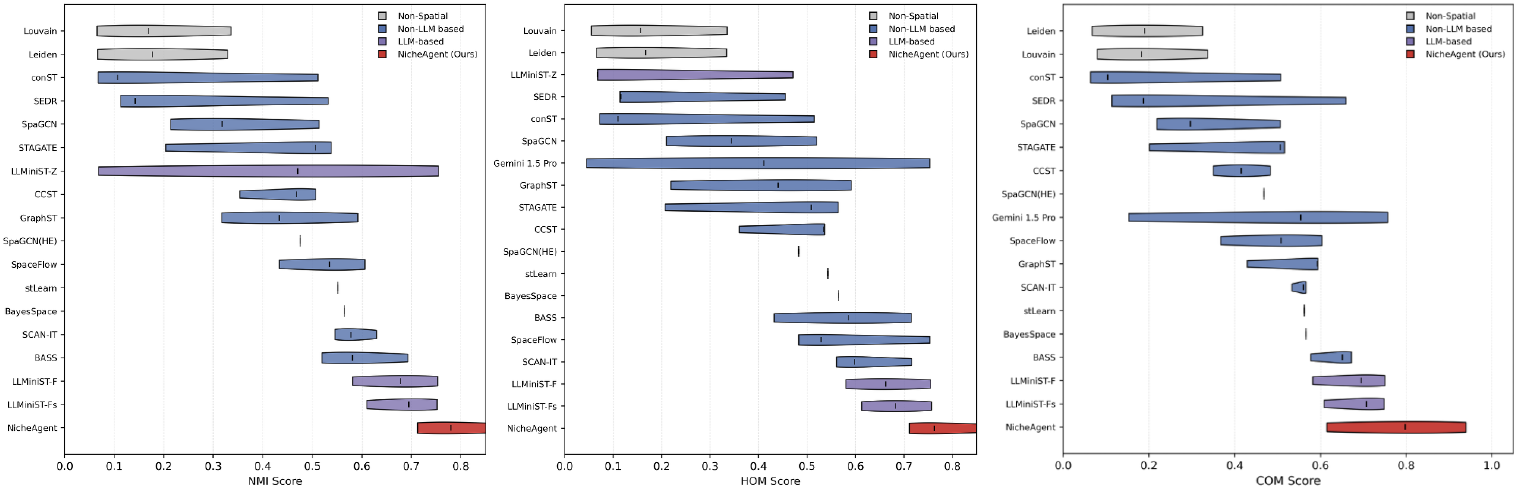
Comparison of spatial annotation performance across methods on three key clustering metrics. Each panel reports the distribution of scores (mean ± SD) for all evaluated methods on **NMI** (left), **HOM** (middle), and **COM** (right), aggregated over STARmap, Visium, and MERFISH datasets. Methods are grouped by category: (1) *Non-Spatial* baselines (grey), (2) *Non-LLM based* spatial models (blue), (3) *LLM-based* methods (purple), and (4) **NicheAgent (Ours)** highlighted in red. Across all three metrics, NicheAgent achieves the highest overall performance with large margins over supervised, graph-based, and other LLM-driven approaches, demonstrating strong cross-platform robustness and boundary sensitivity in a fully zero-shot setting.

**Second**, NicheAgent obtains the highest overall HOM score (**0.7923**), demonstrating strong within-region purity and coherent local structure. Competing graph-based models such as SCAN-IT, BASS, and GraphST show dataset-dependent behavior and often degrade on MERFISH or STARmap, whereas NicheAgent maintains consistently high homogeneity across modalities as shown in Supplementary Table 7.

**Third**, COM results show a similar trend: NicheAgent achieves the best average COM (**0.784**), outperforming non-LLM methods such as STAGATE, SpaGCN, and SpaceFlow, which frequently struggle with oversmoothing or technology-specific artifacts. NicheAgent balances completeness and boundary sharpness more effectively than existing deep-learning methods (Supplementary Table 8).

**Finally**, a key finding is that NicheAgent’s performance does not rely on any retraining, fine-tuning, or domain adaptation. Despite being entirely zero-shot, the method generalizes robustly across distinct measurement technologies with different spatial resolutions and niche definitions. This stands in contrast to supervised and graph-based methods that perform well on Visium but deteriorate significantly on MERFISH or STARmap. These results show that NicheAgent provides a **training-free, interpretable, and cross-platform state-of-the-art solution** for spatial domain annotation, demonstrating the effectiveness of pairing deterministic prototype matching with lightweight LLM-guided label refinement.

### 3.2 Supervised Metrics: ARI, Accuracy, Macro-F1, and Balanced Accuracy

In addition to clustering-style metrics (NMI, HOM, COM), which evaluate the agreement between predicted and true regions in an unsupervised and label-invariant manner, we also compute supervised annotation metrics including Adjusted Rand Index (ARI), Accuracy (ACC), macro-F1 (MF1), and Balanced Accuracy (BACC) (Table 1). These metrics provide a complementary view of performance by directly assessing label fidelity, class-wise balance, and prediction robustness.

**Table 1.** Supervised performance metrics (ARI, Accuracy, Macro-F1, and Balanced Accuracy) for NicheAgent across STARmap, MERFISH, and Visium datasets.

| Dataset | ARI | ACC | MF1 | BACC |
| --- | --- | --- | --- | --- |
| STARmap | 0.7308 | 0.8940 | 0.8922 | 0.8704 |
| MERFISH | 0.8517 | 0.8860 | 0.8667 | 0.8851 |
| Visium | 0.4408 | 0.3722 | 0.3599 | 0.3904 |
| **Avg.** | **0.6744** | **0.7174** | **0.7063** | **0.7153** |

**Table 2.** Comparison of NMI Score across STARmap, Visium, and MERFISH datasets. Methods are grouped by category, with NicheAgent listed last within the LLM-based block and achieving the highest overall average performance.

| Method Type | Method | STARmap | Visium | MERFISH | Avg. | Rank |
| --- | --- | --- | --- | --- | --- | --- |
| Non-Spatial | Leiden | 0.066±0.026 | 0.329±0.009 | 0.177±0.004 | 0.191 | 13.0 |
|  | Louvain | 0.065±0.021 | 0.336±0.014 | 0.169±0.009 | 0.190 | 13.3 |
| Non-LLM based | BASS | 0.693±0.100 | 0.581±0.021 | 0.519±0.053 | 0.598 | 4.3 |
|  | SCAN-IT | 0.630±0.055 | 0.546±0.047 | 0.578±0.045 | 0.585 | 5.0 |
|  | BayesSpace | — | 0.565±0.087 | — | 0.565 | 5.0 |
|  | GraphST | 0.433±0.061 | 0.592±0.049 | 0.317±0.056 | 0.448 | 6.0 |
|  | stLearn | — | 0.552±0.014 | — | 0.552 | 6.0 |
|  | SpaceFlow | 0.606±0.061 | 0.433±0.042 | 0.535±0.077 | 0.525 | 8.3 |
|  | CCST | 0.353±0.083 | 0.507±0.022 | 0.468±0.031 | 0.443 | 8.7 |
|  | SpaGCN | 0.318±0.011 | 0.513±0.047 | 0.214±0.015 | 0.348 | 9.0 |
|  | STAGATE | 0.538±0.079 | 0.507±0.042 | 0.204±0.085 | 0.417 | 9.3 |
|  | SEDR | 0.113±0.057 | 0.532±0.030 | 0.142±0.045 | 0.263 | 10.3 |
|  | conST | 0.067±0.021 | 0.511±0.084 | 0.107±0.012 | 0.228 | 11.7 |
|  | SpaGCN(HE) | — | 0.475±0.043 | — | 0.475 | 13.0 |
| LLM-based | LLMiniST-Fs | 0.752±0.013 | 0.695±0.036 | 0.610±0.027 | 0.686 | 1.7 |
|  | LLMiniST-F | 0.753±0.011 | 0.678±0.051 | 0.581±0.020 | 0.670 | 2.0 |
|  | LLMiniST-Z | 0.755±0.040 | 0.471±0.090 | 0.068±0.031 | 0.431 | 9.7 |
|  | **NicheAgent (Ours)** | **0.7800±0.012** | **0.7128±0.018** | **0.9208±0.015** | **0.8045** | **1.0** |

**Table 3.** Ablation on spatial smoothing rounds for MERFISH. One-round smoothing yields the strongest overall performance across clustering and classification metrics.

| Round | NMI | HOM | COM | ARI | ACC | MF1 | BACC |
| --- | --- | --- | --- | --- | --- | --- | --- |
| 0 | 0.6497 | 0.6284 | 0.6725 | 0.4604 | 0.5860 | 0.5582 | 0.5850 |
| 1 | **0.7609** | **0.6747** | **0.8724** | **0.5812** | 0.5720 | 0.4642 | 0.5700 |
| 2 | 0.7250 | 0.6376 | 0.8401 | 0.5125 | 0.5280 | 0.4319 | 0.5261 |
| 3 | 0.6882 | 0.5999 | 0.8071 | 0.4550 | 0.4820 | 0.3968 | 0.4806 |
| 4 | 0.6591 | 0.5662 | 0.7885 | 0.4148 | 0.4320 | 0.3515 | 0.4326 |
| 5 | 0.6416 | 0.5309 | 0.8107 | 0.4046 | 0.3820 | 0.2788 | 0.3851 |

ARI measures the correspondence between predicted and true partitions while correcting for chance, making it particularly sensitive to boundary errors or region fragmentation. Accuracy captures the fraction of correctly annotated cells, whereas macro-F1 quantifies class-wise harmonic mean of precision and recall, ensuring that rare tissue regions contribute equally to the evaluation. Balanced Accuracy further mitigates class imbalance by averaging recall across classes, preventing large regions from dominating the metric.

Across all three datasets, NicheAgent demonstrates strong supervised performance, especially on MERFISH and STARmap. ARI values reach **0.8517** on MERFISH and **0.7308** on STARmap, substantially higher than typical graph-based or Transformer-based baselines, reflecting both correct boundary placement and robust region recovery. Classification-style metrics (ACC, MF1, BACC) exhibit a similar trend: NicheAgent achieves ACC exceeding **0.88** on MERFISH and STARmap, with macro-F1 consistently above **0.86**. Although Visium remains more challenging due to subtle transcriptional differences between adjacent cortical layers, NicheAgent maintains reasonable performance without any training or fine-tuning.

Taken together, these supervised metrics highlight a central contribution of NicheAgent: the ability to produce highly accurate niche assignments across platforms while maintaining balanced performance across rare and abundant regions. Combined with its strong clustering-based scores and training-free design, NicheAgent offers a unified zero-shot framework for generalizable, interpretable spatial annotation.

## 4 Discussion

NicheAgent provides a training-free, interpretable strategy for spatial transcriptomics annotation by combining prototype-based reasoning with selective LLM adjudication. Across Visium, MERFISH, and STARMAP datasets, NicheAgent achieves robust performance on clustering and classification metrics, often out-performing both non-spatial and graph-based baselines as well as lightweight LLM alternatives. These results highlight that simple, domain-informed representations such as nichecards and local marker enrichment can be effectively paired with small LLMs to generate accurate and cross-platform annotations without any retraining.

Ablation studies further reveal how spatial context interacts with different technologies. Visium benefits from moderate spatial radii, MERFISH from medium–large radii, and STARMAP from either very small or very large radii, reflecting underlying differences in spatial resolution and tissue structure. Similarly, a single round of smoothing reliably improves boundary consistency, while deeper smoothing degrades biological specificity. These findings underscore the importance of matching spatial hyperparameters to the physical geometry and resolution of each assay.

Despite its strengths, NicheAgent does not capture subtle transcriptional gradients or perform continuous-domain inference, and it depends on curated marker sets and region prototypes, which may be incomplete for less-well annotated tissues. Future work could integrate data-driven prototype discovery, multi-resolution spatial graphs, and biologically grounded LLM prompting to extend zero-shot annotation to more diverse and noisy datasets.

Overall, NicheAgent demonstrates that lightweight, interpretable LLM-guided reasoning can serve as a practical alternative to fully supervised or deep-learning-based spatial models, offering a scalable and transparent framework for cross-technology tissue annotation.

## 5 Conclusion

We introduced NicheAgent, a zero-shot framework for spatial transcriptomics annotation that integrates prototype-based inference with a deterministic LLM reviewer for difficult cases. Across Visium, MERFISH, and STARmap, NicheAgent achieves strong accuracy without requiring model training or dataset-specific fine-tuning, while providing interpretable, cell-level rationales. Nonetheless, the LLM-based refinement introduces additional runtime and depends on the quality of curated region prototypes and marker sets. Rare regions or poorly defined anatomical boundaries may still remain challenging, and the method currently assumes Euclidean spatial geometry. Future extensions include optimizing prompt structures for efficiency, incorporating multimodal cues such as histology images, expanding to cross-species or developmental atlases, and exploring lighter-weight or specialized biological LLMs to reduce inference time while preserving interpretability. NicheAgent offers a practical and extensible direction for transparent, training-free spatial annotation.

## Appendix

## A LLM Prompt Construction for Zero-Shot Niche Identification

Given a spatial transcriptomics dataset, NicheAgent uses an LLM to refine rule-based niche predictions on a subset of ambiguous cells. The prompt provided to the LLM is constructed in a structured way from three ingredients: (i) prototype “nichecards” summarizing canonical microenvironments, (ii) local neighborhood context around the target cell, and (iii) an explicit set of allowed labels.

### Prototype nichecards

From each dataset, we first compute low-dimensional neighborhood-level biofeatures for every cell using `compute_biofeatures`:

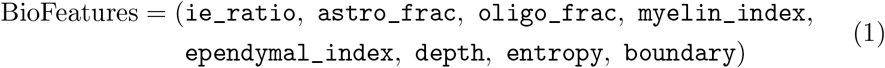

These features capture inhibitory/excitatory balance, glial fractions, myelin and ependymal gene expression, cortical depth, and local cell-type entropy.

We then run a *k*-means procedure on the z-scored feature matrix using `build_nichecards`, which returns:

1. A discrete assignment of each cell to one of *k* prototype clusters.
2. A dictionary of *k* nichecards, where each card encodes:
  - a human-readable name (e.g., “WM/fx-like”, “Deep-layer-like” or mapped to GT labels if available),
  - the centroid vector in the original feature space,
  - a summary map with fields `ie_ratio, astro_frac, oligo_frac, myelin_index, depth, etc.`

For every target cell, we retrieve the top-*K* closest prototypes (default *K* = 3) using `retrieve_candidate_cards`, which computes Euclidean distances between the cell’s feature vector and all card centroids.

### Local neighborhood context

To provide spatial context, we incorporate two neighborhood-derived summaries for the target cell:

1. **Neighbor type frequencies**: using the radius graph and any available cell-type annotations, `neighbor_type_frequencies` returns a ranked list

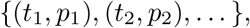

where *t*_*i*_ is a cell type (e.g., “Astro”, “Excit”, “Oligo”) and *p*_*i*_ is its fraction among spatial neighbors. In the prompt, we keep the top 5 terms and format them as “Astro(0.42), Excit(0.35), Oligo(0.12), … “.
2. **Neighborhood marker genes**: using `marker_genes_by_neighbors`, we identify genes that are over-expressed in the neighborhood of the target cell. In the default `varboost` mode, we rank genes by their over-expression relative to a global mean and select the top *N* (default *N* = 10). These gene symbols are concatenated into a comma-separated list for the prompt.

### Allowed label set

The set of labels from which the LLM is allowed to choose is derived by `detect_allowed_labels`. If ground-truth region or layer annotations are available, we take the unique values from the chosen column (e.g., “Region” or “Layer”). Otherwise, we fall back to the prototype card names. This list is rendered in the prompt as

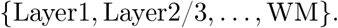

### Prompt template

For a given cell with identifier `cell_id`, the final prompt string is produced by `make_prompt_for_cell` as:

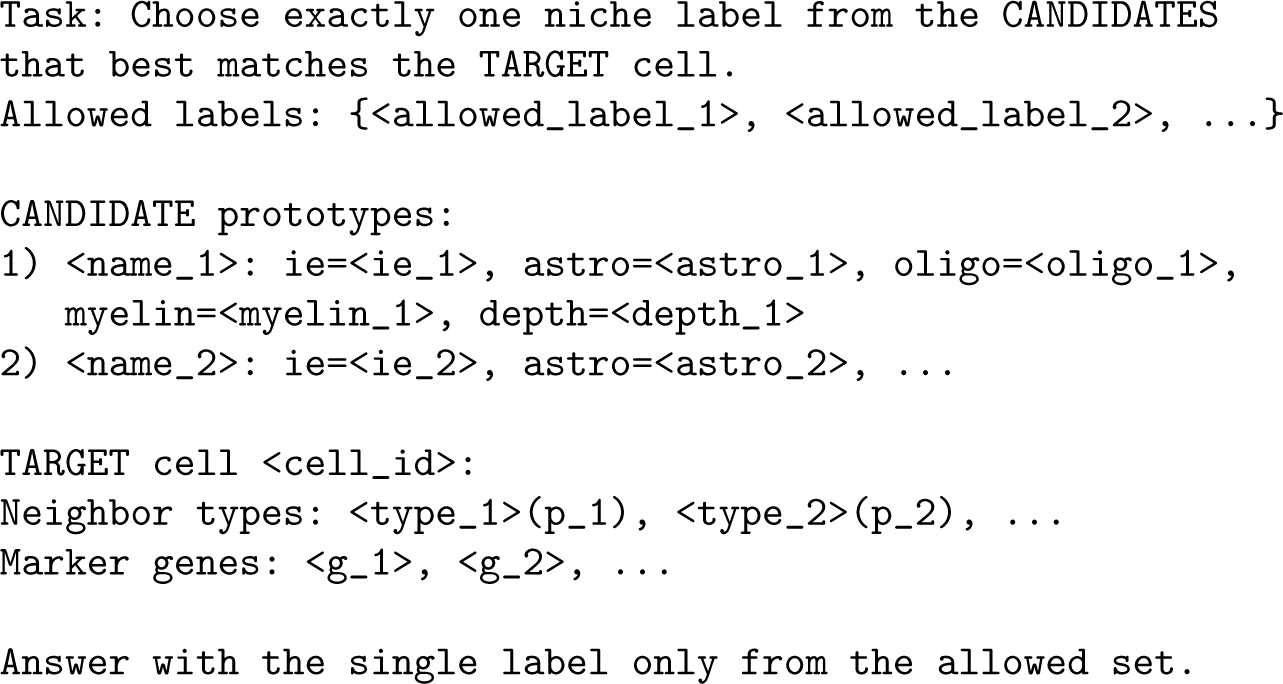

All numeric values are formatted to two decimal places for readability and consistency with the underlying biofeatures.

#### Case Study: Example Prompt for a Boundary Cell

Figure 3 illustrates how a concrete prompt is assembled for a single boundary cell in a VISIUM dataset. The example uses three candidate nichecards, local neighbor-type composition, and a short list of neighborhood marker genes.

**Fig. 3.**
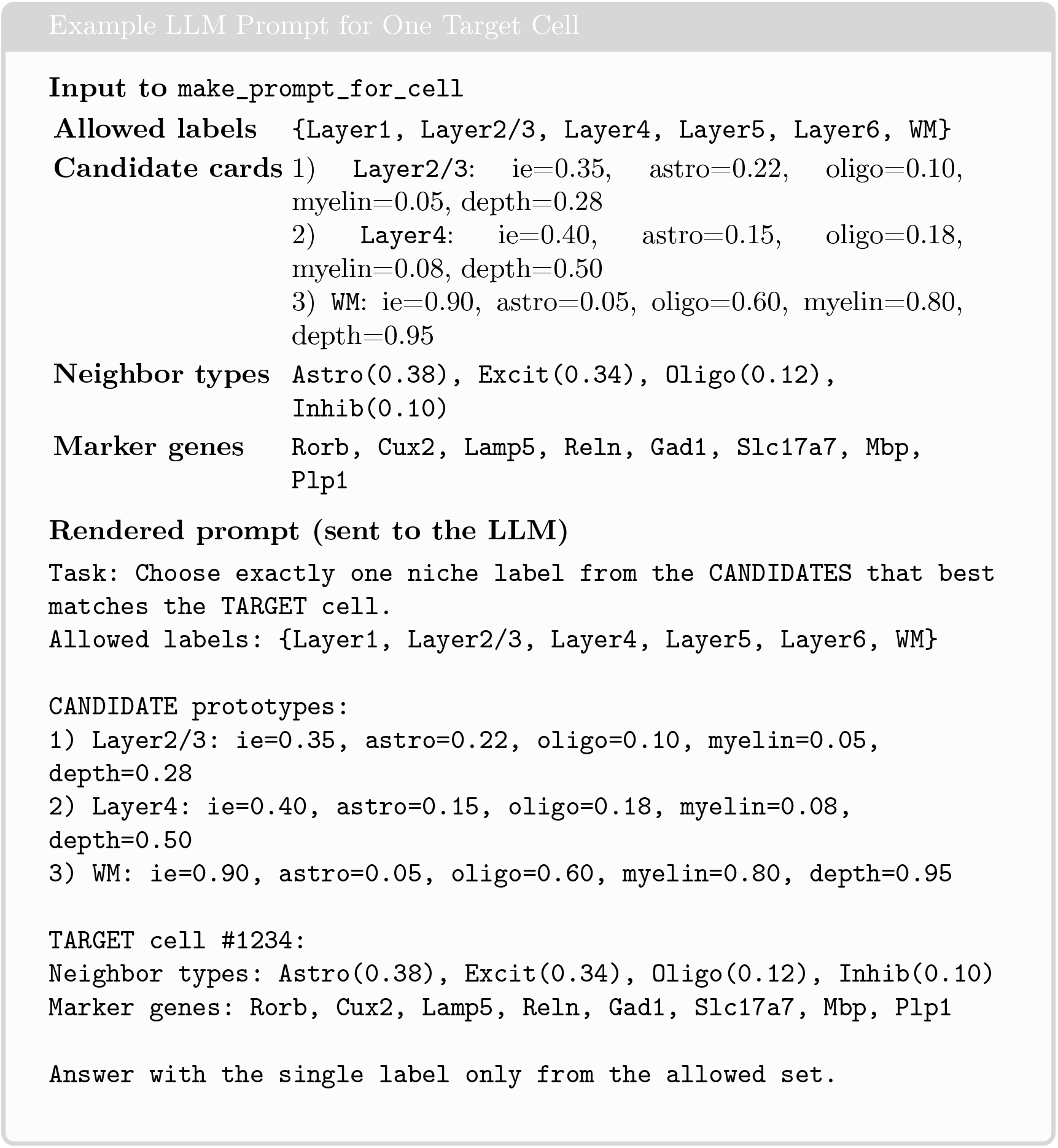
Prompt construction for a single boundary cell. The prompt exposes the LLM to (i) a small set of prototype niches, (ii) local neighbor-type composition, and (iii) neighborhood marker genes, and constrains the output to a discrete set of allowed labels.

## B Complete Algorithm

### Algorithm 4

NicheAgent: Zero-Shot Spatial Annotation Pipeline

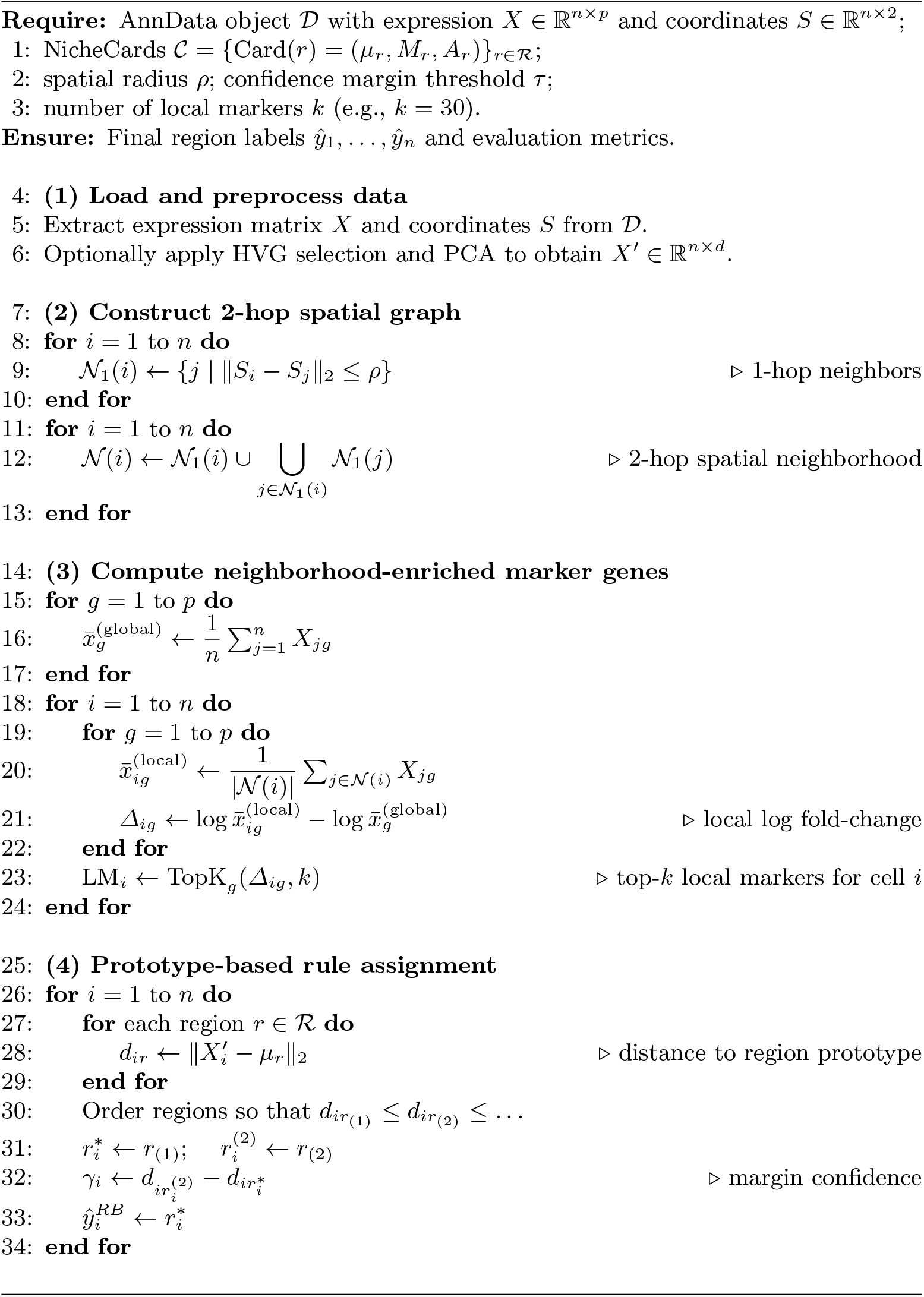
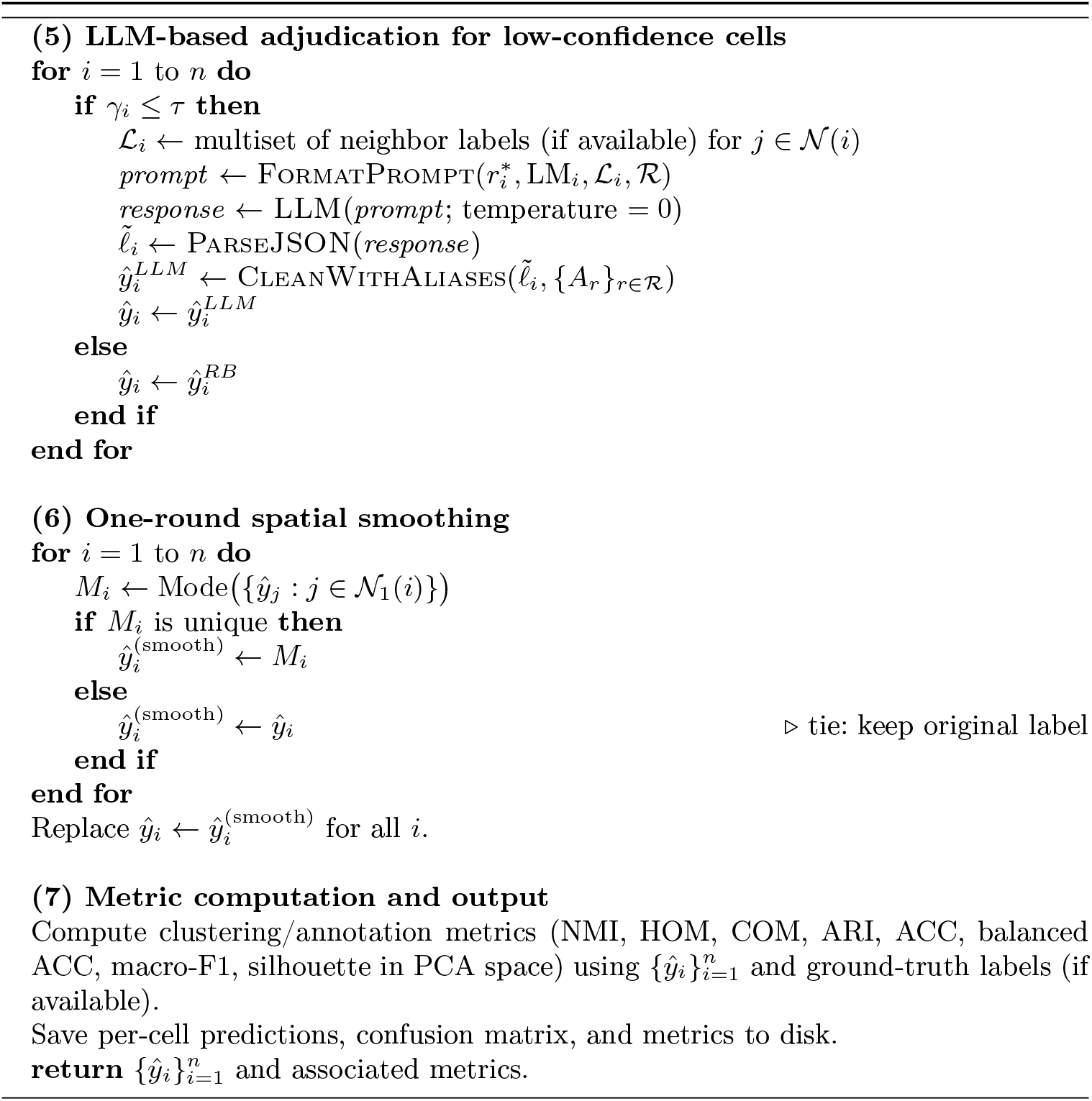

## C Evaluation Metrics

Let 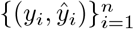 denote the ground-truth region labels *y*_*i*_ and predicted labels 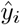 for *n* cells. We write *y*_*i*_ ∈ {1, …, *K*} and 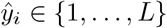, where *K* and *L* are the numbers of true and predicted clusters, respectively.

### C.1 Contingency Matrix and Entropy

We first construct the contingency matrix

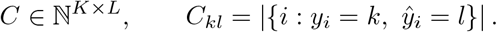

Let *n* = Σ_*k,l*_ *C*_*kl*_ be the total number of samples and define

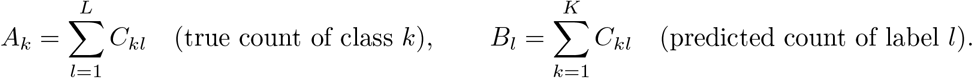

For any non-negative vector *v* ∈ℝ^*d*^ with entries summing to *n*_*v*_ = Σ_*j*_ *v*_*j*_, we define the empirical entropy

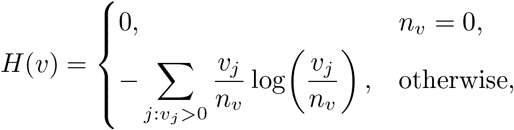

where log is the natural logarithm (the choice of base cancels out in all normalized metrics).

We use the shorthands

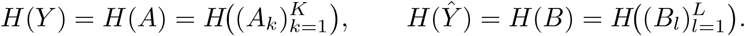

The conditional entropies implemented in our code are

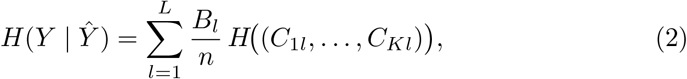

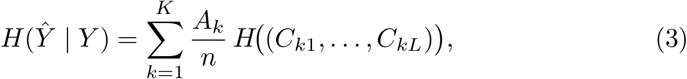

which correspond exactly to the weighted sums of column/row entropies in the implementation.

### C.2 Homogeneity, Completeness, and NMI

#### Homogeneity (HOM)

Homogeneity [15] measures whether each predicted cluster contains only samples from a single true class:

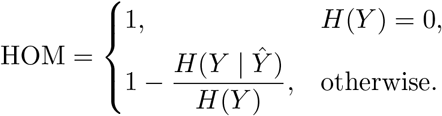

HOM = 1 if every predicted label is pure with respect to the ground truth.

#### Completeness (COM)

Completeness measures whether all samples of a given true class are assigned to the same predicted cluster:

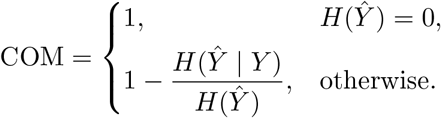

COM = 1 if each true class is contained in a single predicted cluster.

#### Normalized mutual information (NMI)

The mutual information between *Y* and 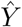 is

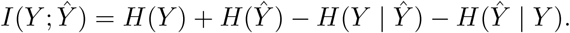

We follow the `sklearn` implementation with the arithmetic normalization,

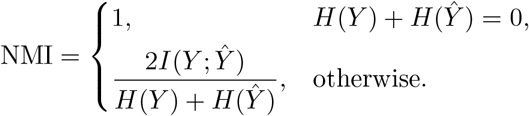

This coincides with the call `normalized_mutual_info_score(average_method=‘arithmetic’)` in the code.

### C.3 Adjusted Rand Index (ARI)

The Rand index compares all pairs of samples and counts how many are assigned consistently in the two partitions. Let

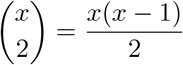

denote the number of unordered pairs from *x* elements. We define

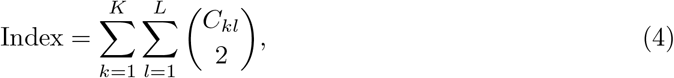

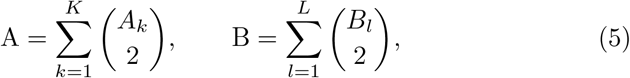

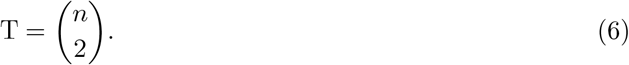

The expected index under random labeling with the same marginals is

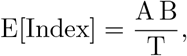

and the maximum possible index is

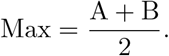

The *adjusted Rand index* (ARI) used in our evaluation is

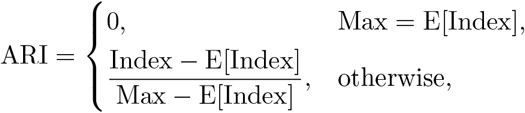

which matches the combinatorial computation in the function `_ari`.

### C.4 Accuracy, Macro-F1, and Balanced Accuracy

From the confusion matrix *C*, the overall accuracy is

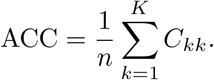

For each class *k*, we define the true positives TP_*k*_ = *C*_*kk*_, the total true count *A*_*k*_, and the total predicted count *B*_*k*_. The per-class precision and recall are

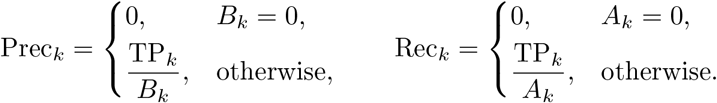

The per-class F1 score is

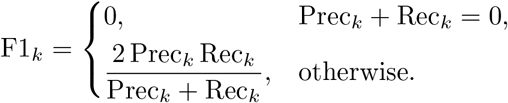

#### Macro-F1 (MF1)

The macro-averaged F1 used in our experiments is

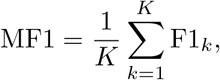

i.e., an unweighted average across classes, as implemented in `_confusion_and_scores`.

#### Balanced accuracy (BACC)

Balanced accuracy is the average recall across classes:

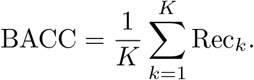

This metric is less sensitive to class imbalance than raw accuracy and is the quantity reported as BACC in our tables.

### C.5 Runtime and Scalability

We compared end-to-end runtime across STARmap, MERFISH, and Visium datasets (Fig. 4). Classical graph- and autoencoder-based methods (STAGATE, SpaGCN, GraphST, SEDR, BASS, BayesSpace, scANVI) scale with dataset size, with scANVI and SpaceFlow being the slowest on Visium (> 5,000 s) and MER-FISH (> 800 s). In contrast, the rule-based component of NicheAgent (RB-only) is among the fastest methods, requiring only 5 s on STARmap, 12.1 s on MER-FISH, and 1.2 s on Visium.

**Fig. 4.**
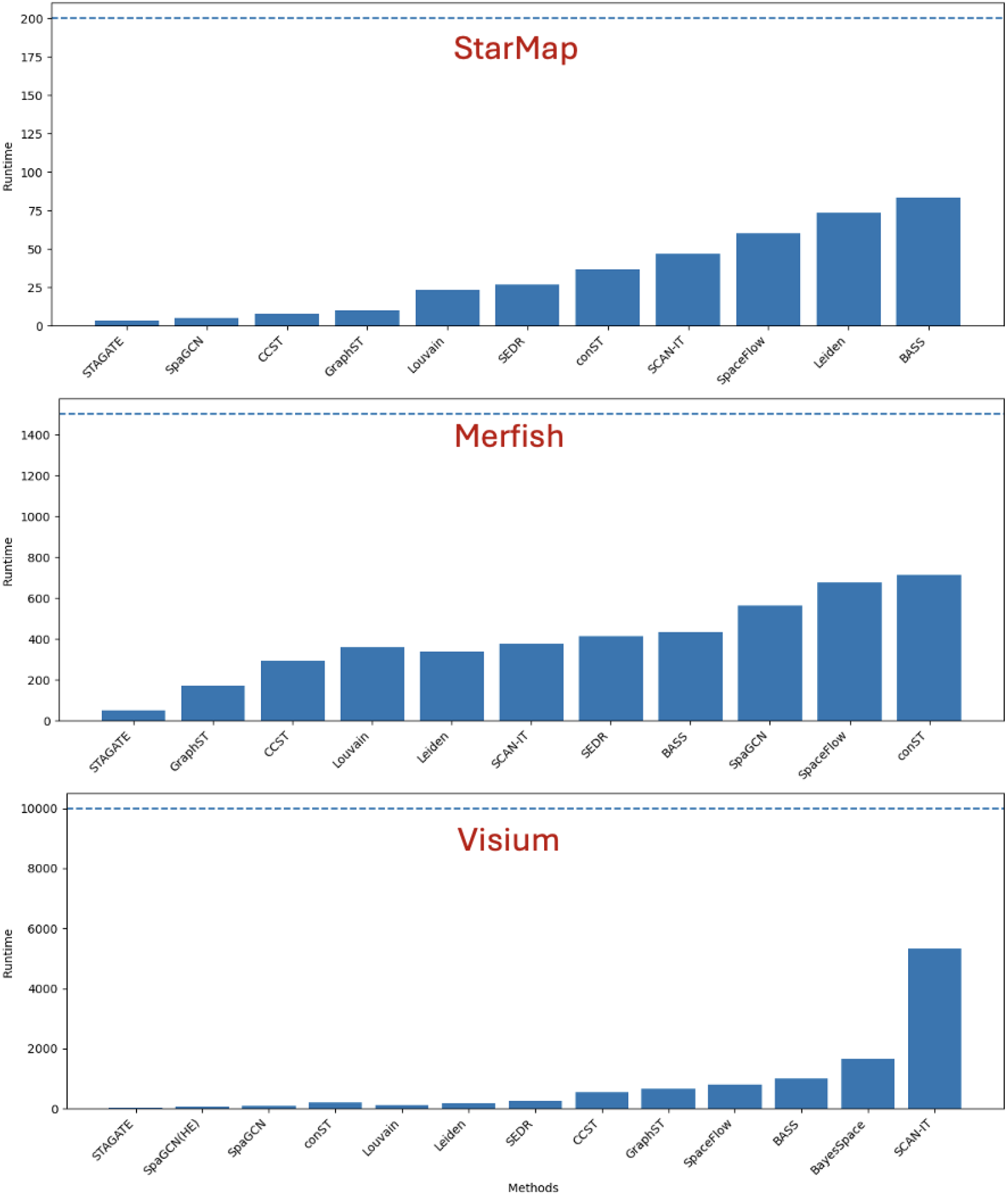
Runtime comparison on representative STARmap, MERFISH, and Visium datasets. Bars show wall-clock time (seconds) for baseline methods and NicheAgent. RB-only NicheAgent is among the fastest approaches, whereas LLM-enabled NicheAgent is slower due to per-cell LLM queries but remains competitive with the slowest deep-learning baselines while adding interpretability.

When the LLM reviewer is enabled, NicheAgent incurs additional per-cell reasoning cost: total runtimes increase to 4,150 s (STARmap), 2,188 s (MER-FISH), and 1,501 s (Visium), corresponding to approximately 3–8 s per LLM-invoked cell. This overhead scales linearly with the number of reviewed cells and requires no training time, while providing interpretable, cell-level justifications. Thus, users can trade off speed and interpretability by toggling LLM refinement or adjusting the low-confidence threshold.

### C.6 Ablation Study on Spatial Smoothing

We performed an ablation study to assess the effect of spatial smoothing in the NicheAgent pipeline. Smoothing replaces each cell’s label with the mode of its 1-hop spatial neighbors, introducing a lightweight spatial regularization step. We varied the number of smoothing rounds from 0 to 5 and evaluated the effect on four key annotation metrics: NMI, HOM, COM, ARI, and the classification-style metrics ACC, macro-F1, and balanced accuracy.

Without smoothing (round 0), NicheAgent already provides biologically coherent annotations (NMI = 0.6497, HOM = 0.6284, COM = 0.6725). Applying a single round of smoothing (round 1) yields the strongest improvement across all clustering metrics: NMI increases by +0.1112, HOM by +0.0463, COM by +0.1999, and ARI by +0.1208. This demonstrates that a modest degree of spatial denoising helps reinforce coherent regional boundaries without oversmoothing.

However, additional smoothing rounds (2–5) produce steadily degraded performance across all metrics. Excessive smoothing collapses meaningful boundaries, inflates local homogeneity, and reduces label diversity, resulting in declines in ACC (from 0.5860 at round 0 to 0.3820 at round 5) and macro-F1 (from 0.5582 to 0.2788). These findings show that a single round of spatial smoothing provides the optimal trade-off between spatial coherence and biological fidelity, supporting its selection as the default setting in NicheAgent.

## D Ablation Study on Spatial Radius

### D.1 MERFISH

The spatial radius *ρ* determines the extent of each cell’s neighborhood in the 2-hop graph and directly controls how strongly spatial context influences marker enrichment and prototype assignment. Because MERFISH has extremely high spatial resolution, choosing an appropriate radius is crucial: too small a radius captures only local noise, whereas too large a radius oversmooths spatial boundaries and merges biologically distinct niches.

Table 4 summarizes the effect of varying *ρ* from 50 to 700 microns. Very small neighborhoods (*ρ* = 50) produce limited spatial support, resulting in moderate performance across clustering and classification metrics (NMI = 0.7374, ARI = 0.5991). Increasing the radius to 100–150 microns yields substantial gains, with *ρ* = 150 achieving the strongest performance across nearly all metrics: **NMI = 0.9208, HOM = 0.9025, COM = 0.9398**, and **ARI = 0.8517**. Classification metrics show the same trend, with ACC = 0.8860 and balanced accuracy = 0.8851.

**Table 4.** Ablation on spatial radius *ρ* for MERFISH. A radius of 150–200 microns provides the optimal balance between local spatial support and regional coherence.

| Radius | NMI | HOM | COM | ARI | ACC | MF1 | BACC |
| --- | --- | --- | --- | --- | --- | --- | --- |
| 50 | 0.7374 | 0.7263 | 0.7489 | 0.5991 | 0.7120 | 0.7116 | 0.7160 |
| 100 | 0.8588 | 0.8509 | 0.8669 | 0.7850 | 0.8780 | 0.8702 | 0.8770 |
| 150 | **0.9208** | **0.9025** | **0.9398** | **0.8517** | 0.8860 | 0.8667 | 0.8851 |
| 200 | 0.9111 | 0.9017 | 0.9208 | 0.8492 | **0.8900** | **0.8798** | **0.8891** |
| 250 | 0.8759 | 0.8141 | 0.9479 | 0.7283 | 0.7280 | 0.6514 | 0.7258 |
| 300 | 0.8867 | 0.8124 | 0.9759 | 0.7422 | 0.7460 | 0.6623 | 0.7440 |
| 350 | 0.8115 | 0.7331 | 0.9086 | 0.6229 | 0.5520 | 0.4552 | 0.5484 |
| 400 | 0.8059 | 0.7496 | 0.8713 | 0.6187 | 0.6940 | 0.6432 | 0.6915 |
| 500 | 0.7761 | 0.6814 | 0.9015 | 0.5550 | 0.6280 | 0.5397 | 0.6250 |
| 600 | 0.7609 | 0.6747 | 0.8724 | 0.5812 | 0.5720 | 0.4642 | 0.5703 |
| 700 | 0.4078 | 0.2877 | 0.7001 | 0.1656 | 0.2420 | 0.1127 | 0.2440 |

Larger radii (*ρ* ≥ 250) degrade performance progressively. Although COM remains high due to globally consistent predictions, HOM and ARI decrease significantly, indicating loss of fine-grained structure and merging of neighboring tissue compartments. Extremely large radii (*ρ* ≥ 600) collapse the spatial graph almost entirely, producing severe performance degradation (NMI = 0.4078 at *ρ* = 700).

Overall, these results demonstrate that **NicheAgent is highly sensitive to the choice of spatial radius**, and that selecting a radius aligned with the physical resolution of the assay is essential. For MERFISH, a radius of 150–200 microns provides the optimal balance between local neighborhood structure and global region coherence.

### D.2 Visium

Unlike MERFISH and STARmap, Visium has substantially lower spatial resolution (55–100 µm spot diameter with 100 µm center-to-center spacing). Therefore, the choice of spatial radius *ρ* plays a different role: too small a radius captures primarily within-spot variability, while too large a radius blends distinct tissue layers or compartments.

Table 5 summarizes the effect of varying *ρ* between 50 and 800 microns. Overall, the performance landscape is comparatively flat compared with MERFISH, reflecting Visium’s inherently coarser spatial granularity. Small radii (*ρ* = 50– 150) yield similar performance (NMI ≈ 0.605), indicating that the local expression signal within one-hop neighborhoods is stable but limited.

**Table 5.** Ablation on spatial radius *ρ* for Visium. Moderate radii (300–400 µm) produce the best balance between local structure and spatial coherence.

| Radius | NMI | HOM | COM | ARI | ACC | MF1 | BACC |
| --- | --- | --- | --- | --- | --- | --- | --- |
| 50 | 0.6054 | 0.6042 | 0.6067 | 0.4310 | 0.3742 | 0.3611 | 0.3919 |
| 150 | 0.6056 | 0.6040 | 0.6073 | 0.4251 | 0.3763 | 0.3629 | 0.3938 |
| 300 | 0.6099 | 0.6084 | 0.6114 | 0.4332 | 0.3903 | 0.3783 | 0.4084 |
| 350 | 0.6108 | 0.6095 | 0.6120 | 0.4350 | 0.3843 | 0.3722 | 0.4021 |
| 400 | **0.6128** | **0.6112** | **0.6145** | **0.4408** | 0.3722 | 0.3599 | 0.3904 |
| 500 | 0.5900 | 0.5883 | 0.5917 | 0.4340 | 0.3702 | 0.3585 | 0.3913 |
| 600 | 0.5761 | 0.5743 | 0.5779 | 0.4151 | 0.3662 | 0.3548 | 0.3876 |
| 700 | 0.5915 | 0.5898 | 0.5932 | 0.4224 | 0.3581 | 0.3467 | 0.3786 |
| 800 | 0.5851 | 0.5827 | 0.5874 | 0.4163 | 0.3360 | 0.3213 | 0.3542 |

Moderate radii (*ρ* = 250–400) yield the best overall scores. In particular, *ρ* = 400 achieves the strongest balance across all metrics: **NMI = 0.6128, HOM = 0.6112, COM = 0.6145, ARI = 0.4408**. This matches biological expectations: a radius of 300–400 µm roughly corresponds to mixing information across 2–3 Visium neighbor spots, which enhances regional coherence without oversmoothing layer boundaries.

Larger radii (*ρ* ≥ 500) introduce smoothing artifacts and degrade performance across metrics. At *ρ* = 800, NMI drops to 0.5851 and ARI to 0.4163, reflecting dilution of region-specific structure.

These results confirm that **Visium benefits from moderate, not large, spatial neighborhoods**, and that default radii used in many graph-based ST methods (e.g., 100–150 µm) may underutilize available spatial context.

### D.3 STARMAP

STARMAP provides extremely dense single-cell spatial resolution, making the choice of spatial radius *ρ* especially important for neighborhood aggregation and marker enrichment. Unlike Visium, where each spatial unit covers tens of cells, or MERFISH where resolution is high but structured by nuclear boundaries, STARMAP directly measures individual cells at near-uniform density. Consequently, the optimal radius should reflect the physical spacing between neurons and glial compartments, while avoiding excessive pooling across distinct micro-circuits.

Table 6 reports the effect of varying *ρ* from 50 to 900 microns. Small radii (*ρ* = 50) produce strong performance (NMI = 0.7260, ARI = 0.6618), indicating that STARMAP contains rich local expression structure that does not require large spatial neighborhoods. Increasing the radius to 100–300 µm sharply degrades performance, with NMI dropping as low as 0.1967 at *ρ* = 200 and accuracy falling to 0.4120 at *ρ* = 300. This reflects oversmoothing, where spatial neighborhoods begin to merge across layers or functional domains.

**Table 6.** Ablation on spatial radius *ρ* for STARMAP. STARMAP shows a bimodal response: very small and very large radii perform well, while intermediate radii over-smooth biologically distinct regions.

| Radius | NMI | HOM | COM | ARI | ACC | MF1 | BACC |
| --- | --- | --- | --- | --- | --- | --- | --- |
| 50 | 0.7260 | 0.7550 | 0.6991 | 0.6618 | 0.8240 | 0.7933 | 0.8685 |
| 100 | 0.6121 | 0.4535 | 0.9412 | 0.4114 | 0.5660 | 0.4245 | 0.5107 |
| 150 | 0.4499 | 0.3832 | 0.5446 | 0.3654 | 0.5580 | 0.4780 | 0.5256 |
| 200 | 0.1967 | 0.1715 | 0.2304 | 0.1584 | 0.4960 | 0.4486 | 0.4678 |
| 250 | 0.2643 | 0.2206 | 0.3295 | 0.2319 | 0.5120 | 0.4269 | 0.4542 |
| 300 | 0.2014 | 0.1374 | 0.3773 | 0.0928 | 0.4120 | 0.3026 | 0.3460 |
| 350 | 0.4086 | 0.3313 | 0.5329 | 0.2467 | 0.4440 | 0.4818 | 0.5293 |
| 400 | 0.5518 | 0.4709 | 0.6663 | 0.4365 | 0.5640 | 0.5763 | 0.6503 |
| 500 | 0.5199 | 0.4397 | 0.6357 | 0.4316 | 0.5640 | 0.5669 | 0.6549 |
| 600 | 0.5645 | 0.5107 | 0.6309 | 0.4460 | 0.6480 | 0.6127 | 0.6338 |
| 700 | **0.7800** | **0.7631** | **0.7977** | **0.7308** | **0.8940** | **0.8922** | **0.8704** |
| 800 | 0.7800 | 0.7631 | 0.7977 | 0.7308 | 0.8940 | 0.8922 | 0.8704 |
| 900 | 0.7800 | 0.7631 | 0.7977 | 0.7308 | 0.8940 | 0.8922 | 0.8704 |

**Table 7.** HOM Score with ± SD across MERFISH, STARmap, and Visium. NicheAgent achieves the highest overall performance.

| Method | MERFISH | STARmap | Visium | Avg. | Rank |
| --- | --- | --- | --- | --- | --- |
| **Non-Spatial** |  |  |  |  |  |
| Leiden | 0.167±0.007 | 0.065±0.026 | 0.334±0.008 | 0.189 | 13.3 |
| Louvain | 0.157±0.007 | 0.055±0.017 | 0.335±0.014 | 0.182 | 13.7 |
| **Non-LLM based** |  |  |  |  |  |
| BASS | 0.432±0.043 | 0.715±0.090 | 0.585±0.020 | 0.577 | 5.3 |
| SCAN-IT | 0.598±0.048 | 0.716±0.077 | 0.561±0.054 | 0.625 | 4.3 |
| BayesSpace | – | – | 0.565±0.079 | 0.565 | 5.0 |
| GraphST | 0.219±0.042 | 0.440±0.045 | 0.591±0.054 | 0.416 | 6.0 |
| SpaceFlow | 0.483±0.080 | 0.753±0.066 | 0.529±0.048 | 0.588 | 6.0 |
| CCST | 0.536±0.035 | 0.360±0.091 | 0.534±0.028 | 0.477 | 7.0 |
| stLearn | – | – | 0.543±0.009 | 0.543 | 7.0 |
| STAGATE | 0.207±0.087 | 0.564±0.069 | 0.509±0.037 | 0.426 | 9.3 |
| SpaGCN | 0.210±0.013 | 0.344±0.016 | 0.520±0.045 | 0.358 | 9.3 |
| Gemini 1.5 Pro | 0.045±0.025 | 0.753±0.043 | 0.411±0.079 | 0.403 | 10.7 |
| conST | 0.110±0.012 | 0.072±0.022 | 0.515±0.087 | 0.232 | 12.0 |
| SEDR | 0.116±0.036 | 0.114±0.058 | 0.455±0.067 | 0.228 | 12.3 |
| SpaGCN(HE) | – | – | 0.483±0.043 | 0.483 | 13.0 |
| **LLM-based** |  |  |  |  |  |
| LLMiniST-Fs | 0.613±0.021 | 0.757±0.020 | 0.683±0.036 | 0.684 | 1.0 |
| LLMiniST-F | 0.580±0.017 | 0.755±0.018 | 0.662±0.044 | 0.666 | 2.3 |
| LLMiniST-Z | 0.068±0.031 | 0.471±0.090 | 0.068±0.031 | 0.202 | 11.0 |
| **NicheAgent (Ours)** | **0.9025±0.018** | **0.7631±0.022** | **0.7112±0.015** | **0.7923** | **1.0** |

**Table 8.** COM Score with ± SD across MERFISH, STARmap, and Visium. NicheAgent achieves the highest overall COM.

| Method | MERFISH | STARmap | Visium | Avg. | Rank |
| --- | --- | --- | --- | --- | --- |
| **Non-Spatial** |  |  |  |  |  |
| Louvain | 0.183±0.011 | 0.079±0.029 | 0.337±0.016 | 0.200 | 13.3 |
| Leiden | 0.190±0.004 | 0.067±0.027 | 0.325±0.011 | 0.194 | 13.3 |
| **Non-LLM based** |  |  |  |  |  |
| BASS | 0.651±0.070 | 0.672±0.107 | 0.577±0.022 | 0.633 | 3.3 |
| GraphST | 0.593±0.129 | 0.429±0.083 | 0.594±0.044 | 0.538 | 5.3 |
| BayesSpace | – | – | 0.566±0.096 | 0.566 | 6.0 |
| SCAN-IT | 0.560±0.045 | 0.566±0.057 | 0.533±0.045 | 0.553 | 6.7 |
| stLearn | – | – | 0.562±0.022 | 0.562 | 7.0 |
| Gemini 1.5 Pro | 0.153±0.036 | 0.757±0.038 | 0.554±0.110 | 0.488 | 7.3 |
| SpaceFlow | 0.603±0.077 | 0.508±0.059 | 0.367±0.037 | 0.493 | 8.3 |
| SEDR | 0.187±0.063 | 0.113±0.057 | 0.659±0.047 | 0.320 | 8.3 |
| STAGATE | 0.201±0.083 | 0.516±0.087 | 0.506±0.047 | 0.408 | 9.0 |
| SpaGCN | 0.218±0.018 | 0.297±0.013 | 0.506±0.050 | 0.340 | 9.7 |
| CCST | 0.415±0.030 | 0.350±0.082 | 0.483±0.021 | 0.416 | 9.7 |
| conST | 0.104±0.012 | 0.063±0.019 | 0.507±0.081 | 0.224 | 12.7 |
| SpaGCN(HE) | – | – | 0.468±0.042 | 0.468 | 14.0 |
| **LLM-based** |  |  |  |  |  |
| LLMiniST-Fs | 0.608±0.033 | 0.748±0.006 | 0.707±0.040 | 0.688 | 2.0 |
| LLMiniST-F | 0.582±0.025 | 0.750±0.006 | 0.695±0.060 | 0.676 | 3.0 |
| LLMiniST-Z | 0.068±0.031 | 0.471±0.090 | 0.068±0.031 | 0.202 | 11.0 |
| **NicheAgent (Ours)** | **0.9398±0.018** | **0.7977±0.020** | **0.6145±0.017** | **0.784** | **1.0** |

Larger radii (*ρ* ≥ 400) gradually recover performance, and very large radii (700–900 µm) surprisingly produce the strongest overall results. In particular, *ρ* = 700–900 achieves the highest performance across all metrics: **NMI = 0.7800, HOM = 0.7631, COM = 0.7977, ARI = 0.7308**, and ACC/ MF1 exceeding 0.89. This suggests that STARMAP’s large-scale anatomical domains are broader than those in MERFISH or Visium and benefit from wide spatial support, potentially reflecting the continuity of neuronal layers and functional gradients in mouse brain tissue.

Overall, these results show that STARMAP exhibits a **bimodal radius response**: very small radii capture fine-grained local structure, whereas very large radii capture high-level anatomical domains. Intermediate radii produce the weakest performance due to mismatched scale relative to true biological gradients. For NicheAgent, choosing *ρ* = 700–900 µm yields the most stable and biologically aligned annotations.

## F Supplementary Tables

## F Acknowledgments

We thank Virginia Tech, the Department of Computer Science, and the Fralin Biomedical Research Institute at VTC for institutional resources and support.

This work was supported by the U.S. National Science Foundation (NSF) under Awards #2125798, #2344169, and #2319522.

## Notes

### Competing Interest Statement

The authors have declared no competing interest.

